# miR-1290 stimulates proliferation of gastric cancer by targeting SP1 with overexpression of fucosyltransferase IV and α1, 3-fucosylated glycans

**DOI:** 10.1101/2020.01.05.895300

**Authors:** Faisal Aziz, Li Yulin, Qiu Yan

**Affiliations:** Cellular and Molecular Biology Division, The Hormel institute-University of Minnesota, Austin, MN, USA; Department of Biochemistry and Molecular Biology, Dalian Medical University, Liaoning Provincial Core Lab of Glycobiology and Glycoengineering, Dalian, PRC

**Keywords:** Fucosyltransferase IV, gastric cancer, *Helicobacter pylori*, miRNA-1290, SP1, ubiquitination

## Abstract

Fucosylation plays an important role in the development of carcinogenesis. miRNA-1290 emerged as crucial molecule to regulate cancer cell proliferation. This study evaluated the role of miRNA-1290 to development of gastric cancer by regulation of fucosyltransferase-IV, specific protein-1 (SP1) and α1,3-fucosylated glycans.We analyzed the role of *H. pylori* and miR-1290 in gastric cancer cells in induce fucosylation and cell proliferation, as well as SP1 and ubiquitin protein interaction. We found miR-1290 induced proliferation in *H. pylori* CagA treated gastric cancer cells by stimulating FUT4/LeY fucosylation, as evidence by high expression of miR-1290 and phosphorylation of EGFR and MAPKs pathway in dose–dependent manner. In addition, miR-1290 inhibited SP1 protein with the regulation of ubiquitin-proteasomal system and leads to stimulate FUT4 and α1,3-fucosylated glycans level. We report the role of miRNA-1290 to stimulate FUT4 fucosylation and LeY through EGFR/MAPKs pathway by targeting SP1 in the development of gastric cancer.

## Introduction

Gastric cancer is the second most common cancer in cancer-related deaths. It accounts 42% of all gastric cancer cases worldwide [1], [2]. *Helicobacter pylori* (*H. pylor*i) colonization is associated with chronic gastritis, gastric ulcer and gastric cancer [3], [4], which colonizes 50% of the humans’ stomach [5], [6]. The International research agency for cancer (IRAC) ranked *H. pylori* as a class I carcinogen [7]. CagA toxin (first bacterial oncoprotein) is one of the most important virulence factor produced by *H. pylori* [8], associated with gastric cancer by inducing inflammation, cell proliferation and metastasis [9], [10].

Glycosylation is essential in many of the physiological and pathological processes. *H. pylori* infection alters the host’s glycosylation with the stimulation of specific glycosyltransferases and the sugar antigens such as Lewis blood group antigens [11], [12], [3]. Cancers are characterized by altered fucosylation where fucose units are transferred to precursor oligosaccharides by the catalyzation of fucosyltransferases (FUTs) [13], [14]. High expression of FUT4 promotes cell proliferation through augmenting the synthesis of LeY. LeY is difucosylated oligosaccharide that belongs to A, B, H and Lewis blood group family with the chemical structure [Fucα1→2Galβ1→4(Fucα1→3) GlcNAcβ1→R]. The α1, 3-fucosylation of LeY is catalyzed by FUT4 [15]. The role of FUT4 fucosylation and α1,3-fucosylated glycans in the development and progression of *H. pylori* associated gastric cancer has not been well studied.

MicroRNAs (miRNAs) are small non-coding RNA sequences of about 22 nucleotides that negatively regulate protein expression at the posttranscriptional level [16], [17], [18]; by binding to their 3′ untranslated region (UTR) of target mRNAs [19], [20]. miRNA involvement in the *H. pylori* infectious and gastric cancer has been studied [21], [22], [23], [24] but there is need to study the mechanism that how *H. pylori* CagA contributes to promote gastric cell proliferation with the activation of miRNA. miRNA-1290 is targeted and altered several potential transcription factors genes [25], including Foxa1, Smad2, Bach1, MITF, and HoxC13 [26]. FUT4 is regulated by various transcriptional and post-transcriptional factors. SP1 is a well-characterized DNA-binding protein which augments cell growth and metastatic potential of tumor cells [27]. High expression of SP1 is inversely proportional to cancer development and progression [28], [29]; and associated with poor prognosis in human cancers such as breast [30], colon [31], lung [32] and gastric cancer [33], [34].

We previously reported that FUT4 expression is downregulated by SP1 and induce gastric cancer apoptosis [34], [35]. The role of miRNA-1290 in regulating SP1 transcription factors in gastric cancer has not been studied. We hypothesized that SP1 might be potentially specific target of miRNA-1290 with the overexpression of FUT4/LeY through EGFR/MAPKs pathway for the development of gastric cancer, which might be a new pathogenic mechanism of *H. pylori* CagA-associated gastric cancer.

## Material and Methods

### Bacterial cultivation

*H. pylori* strain (ATCC-43504) was processed and cultured on Colombia agar (CA) plates containing 7% lysed horse blood and antibiotics (Amphotericin B, Trimethoprim, Cefsulodin and Vancomycin). The CA plates were incubated for 3–4 days under microaerophilic conditions at 37 °C, and identification was carried out by different conventional and molecular methods.

### Cytotoxin-associated immunodominant antigen-pathogenicity island protein (Cag PAI) with type IV system

Cytotoxin-associated immunodominant antigen-pathogenicity island protein (Cag-PAI) encoded by the cagA (cag26)(cai)(HP_05470) gene of *H. pylori*. Codon optimized cDNA sequence was designed for recombinant protein production in both *E. coli* and mammalian cell (Bioclone, US) with protein

### Cell culture

SGC-7901 gastric cancer epithelial cell line (Shanghai Institute of Cell Biology, Shanghai, China) were grown in Dulbecco’s Modified Eagle’s Medium (DMEM)/F12 supplemented with 10% FBS (Invitrogen, Camarillo, CA, US)., 5 µg/ml insulin, 100 U/ml penicillin and 100 µg/ml streptomycin; maintained at 37 °C under 5% CO_2_. Cells were seeded in 100 mm dish of DMEM/F12. When the cells reached to 70% confluence, cells were washed three times with phosphate-buffered saline (PBS) and infected with *H. pylori* (1×10^3^ bacteria/ml) as well as treated with different concentrations of *H. pylori* CagA PAI as 5, 10 and 15 µg/ml under serum starved conditions [36].

### Transient transfection

Human gastric cancer cells (SGC-7901) were trypsinized and seeded into 6-well plates. When cells reached to 70–80% confluence, FUT4 siRNA (Zhang et al. 2008) was transiently transfected into SGC-7901 cells using 400 ng of plasmid and 2 µl of Lipofectamine™ Reagent and Plus™ Reagent (Invitrogen) according to manufacturer’s instructions. The transfection reagent was removed after 12 h and was followed by CagA treatment. Cells were harvested 24 and 48 h post-CagA treatment.

### Knockdown and overexpression of miRNA-1290

Cells were plated at 1.5−2×10^3^ and 1×10^5^ cells per well in 96- and 6-well plates one day before transfection, and at 70% confluency transfected with Dharmacon hsa-miRNA-1290 mimic or hairpin inhibitor with appropriate control of miRNAs at 30nM final concentrations using DharmaFECT-2 transfection reagent according to manufacturer’s instructions.

### Enzyme-linked immunosorbent assay

Serological assessment of FUT4 and LeY were measured by enzyme-linked immunosorbent assay (ELISA). Commercial ELISA kits were used to analyze serological values of FUT4 according to manufacturer’s instructions (Cusabio, US). LeY serological assessment was detected by in-house ELISA as described [3], [34].

### Cell proliferation and viability assay

Cells (1 × 10^3^ cells/well) were plated in 96-well plates. After 24 h, cells were treated with the *H. pylori*, as well as CagA protein, incubated at 37 °C for 24, 48 or 72 h. CCK8 (Dojindo CK40, Japan) was used to detect cell proliferation. Moreover, cells were treated with different inhibitors, including pEGFR (AG-1478), ERK (PD-98059), p-38 (SB-203580), SP1 (Mithramycin A), miRNA-1290 and FUT4 SiRNA. In brief, 10 µl CCK8 was added to the culture medium and incubation was continued for 4 h at 37 °C. Absorbance was measured at 450 nm. The cell viability assay was also performed to determine the viability of gastric cancer cells by trypan blue exclusion assay. The test was performed in six replicates.

### Semi-quantitative RT-PCR and real-time PCR

Total RNA was extracted by using the TRIzol® reagent (Invitrogen) and synthesized (Takara). The real-time PCR (qRT-PCR) was carried out according to the manufacturer’s protocol (Takara) (Supplementary table1).

### Immunoprecipitation assay

Immunoprecipitation assay was used to analyses the interaction between SP1 and ubiquitin. The SGC-7901 cells were treated with *H. pylori* CagA and harvested (3 × 10^6^) in lysis buffer [150 mM Tris-HCl (pH 7.5), 1M NaCl, 0.5M EDTA (pH 8.0), 0.5M EGTA (pH 7.5), 2 mM Na3 VO4, 1M NaF, 20% SDS, Glycerol 50%, and Triton] with continuous circular rocking for 30 min at 4°C. The lysates were centrifuged at 12,000 rpm for 20 min at 4°C and concentration of supernatant fraction was measured using BCA protein Assay kit. Total cell lysates (300 μg) supernatant fractions were used for immunoprecipitation with their respected primary antibody (1 μg) of SP1 and ubiquitin and 20μl glutathione-Sepharose or protein A. incubated for overnight at 4°C with continuous rocking shaking. Later, the complexes were collected by centrifugation, washing with lyses buffer and remove the supernatant. Reactions were stopped with 6X SDS buffer, separated by 12% SDS-PAGE, and visualized by Western blot.

### Anchorage-independent cell growth assay

SGC7901 cells (8 × 10^3^/well) were suspended in 1 mL basal medium Eagle (BME) medium containing 10% FBS and 0.33% agar with the indicated concentrations of different inhibitors, including EGFR (AG-1478), ERK (PD-98059), p-38 (SB-203580), SP1 (Mithramycin A), miRNA-1290, and FUT4 SiRNA, plated on 3 ml of solidified BME containing 10% FBS and 0.5% agar with the same indicated concentrations of different inhibitors as on the top layer. The cultures were maintained at 37°C in a 5% CO_2_ incubator for 1–4 weeks. Colonies were counted under a microscope using (Nikon Ti-DS, Japan). Cells without any treatment or treated with CagA protein treated as control. The assay was performed in triplicates.

### Indirect immunofluorescent & immunocytochemical staining

Cells were grown on glass coverslips. After 48 h of CagA treatment, evaluated for rabbit anti-PCNA (1:100) and mouse anti-p-EGFR (1:100) as described [3], [34].

### Immunoblotting

Cells were washed with PBS (pH7.4), and incubated with lysis buffer and processed and evaluated for anti-PCNA (1:1000), anti-SP1 (1:1000) anti-FUT4 (1:1000); Proteintech Group, anti-LeY (1:1000), anti-LeA (1:1000), anti-LeB (1:1000), anti-LeX (1:1000); Abcam (Cambridge, USA), anti-sLeX (KM93, IgM) (1:500); Millipore, USA., anti-Ubiquitin (1:500) (Millipore) anti-EGFR (1:1000) and anti-pEGFR (1:500), anti-ERK1/2 (1:500) and anti-pERK1/2 (1:500), anti-p38 (1:1000) and anti-pp38 (1:1000); (Transduction Laboratories, US), as described [3], [34].

### Flow cytometry assay

Cells were gently trypsinized to prepare a single-cell suspension and were incubated with anti-PCNA (1:100), anti-p-EGFR (1:100), anti-FUT4 (1:100), and anti-LeY (1:100) as described [3], [34].

### Statistical analysis

Each experiment was repeated 3 times independently and results presented as the mean ± SEM. Statistical differences between test groups were analysed by paired Student’s *t-*test and one-way analysis of variance (ANOVA) followed by (post hoc) Newman-Keuls multiple comparison tests. Linearity of correlation was calculated by the Pearson’s coefficient correlation method. *P*<0.05 and *P*<0.01 were considered as significant. The statistical software GraphPad prism 5.03 was used for analyzing the data.

## Results

### H. pylori CagA induces gastric cancer cell proliferation with the regulation of EGFR/SP1 through miRNA-1290 overexpression

The role of *H. pylori* CagA was examined to determine the role of miRNA-1290 in the proliferation of gastric cancer cells. As shown in Figure 1, gastric cancer cell proliferation was significantly increased in CagA treated cells in a dose-dependent manner with the phosphorylation of EGFR by western blot and cytometry assay (*P*<0.001) (Figures. 1A, 1B, 1C and 1I; Supplementary Fig. S1). Colony formation and CCK8 cell proliferation assay were further confirmed significant stimulatory effect of *H. pylori* CagA (*P*<0.001) (Figures. 1D and Supplementary Fig. S1). miRNA-1290 was found to be a putative target of miRNA for *H. pylori* CagA in gastric cancer as miRNA-1290 was significantly stimulated in *H. pylori* CagA treated cancer cells (*P*<0.001) (Figure. 1E). Moreover, *H. pylori* CagA is capable of regulating SP1 transcription factor expressions in gastric cancer cells. As shown in Figure 1, we found that SP1 expression was significantly decreased in *H. pylori* CagA treated cells as compared to untreated cells (*P*<0.001). Furthermore, to identify the mechanism involved in *H. pylori* CagA associated up-regulation of p-EGFR, cells were pretreated with EGFR inhibitor (Tyrphostin AG-1478, 10^−4^ M) followed by *H. pylori* CagA treatment, resulted in the significant inhibition of CagA-activated EGFR phosphorylation, FUT4, miRNA-1290 and gastric cancer cell proliferation while SP1 stimulated as compared to untreated cells (*P*<0.001) (Figures 2F, 2G). Furthermore, Colony formation assay showed the consecutive result as significant stimulatory effect of *H. pylori* CagA on EGFR phosphorylation associated cell proliferation, while inhibited in EGFR inhibitor (AG-1478) treated cells (*P*<0.001) (Fig. 2H). These results predict the likely involvement of p-EGFR associated miRNA-1290 activation in the gastric cancer cell proliferation with the regulation of SP1/FUT4/LeY.

**Figure 1.**
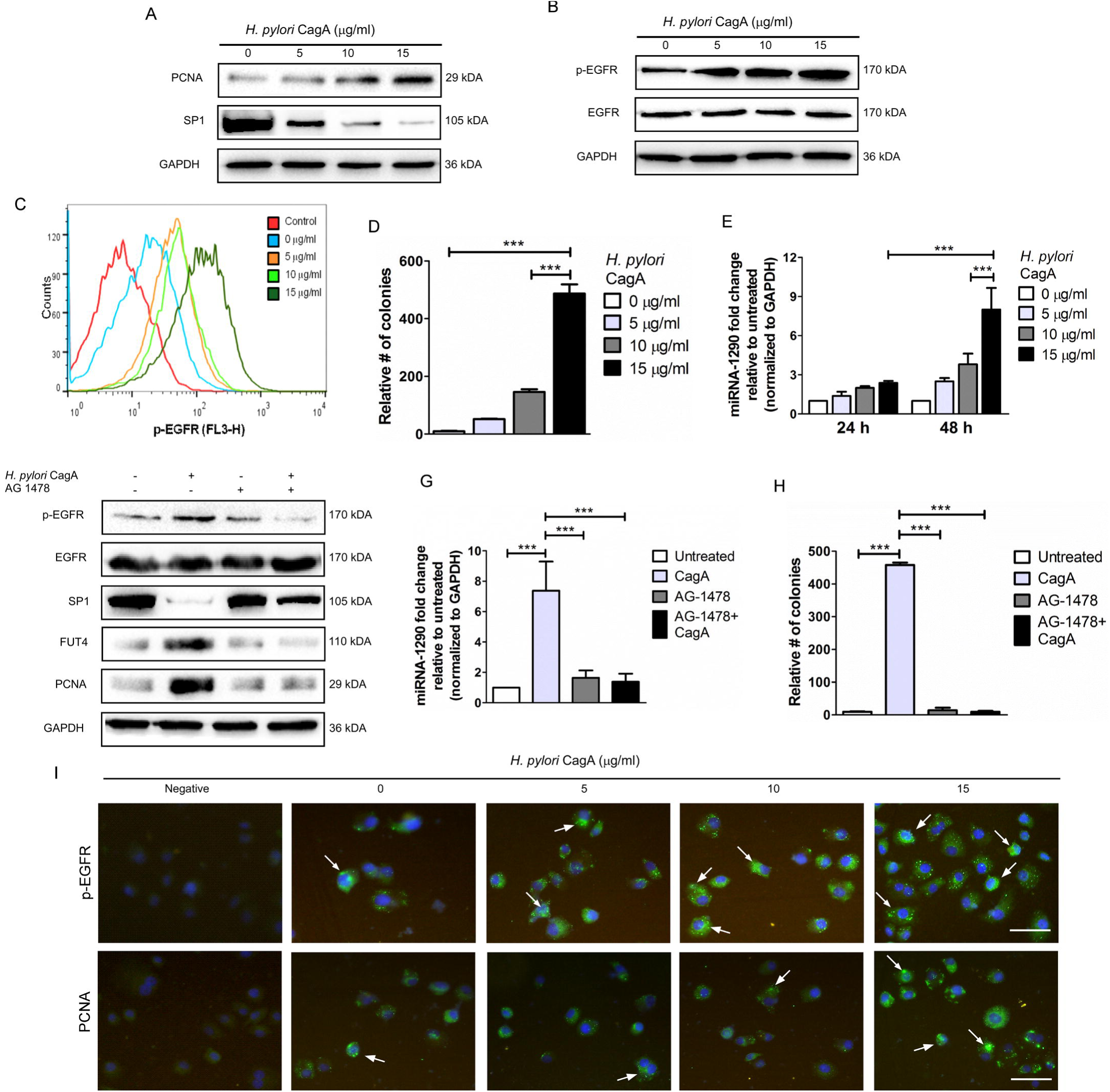
*H. pylori* CagA promotes the cell proliferation with the activation of human gastric cancer cells (SGC-7901). Cells (1 × 10^3^ cells/well) were seeded in 96-well plate and followed by the CagA treatment. The expression of (A) PCNA and (B) EGFR was detected of post CagA treatment by (A and B) Western blot (C) flow cytometry assay and (I) immunofluorescence. Cell proliferation was determined by (D) MTT and (E) anchorage-independent cell growth assay. (E) Relative miRNA-1290 expression in dose-dependent manner after *H. pylori* infection by real time PCR. Moreover, (F) human gastric cancer cells were treated with Tyrphostin AG 1478 (10^−4^M) for 24 h in serum free medium followed by CagA, total proteins from the whole cell lysate were analysed for p-EGFR, SP1, FUT4 and PCNA by Western blotting. (E) (F) Relative miRNA-129 expression was determined by real time PCR after treatment of *H. pylori* CagA or EGFR inhibitor. Cells (1 × 10^3^ cells/well) were seeded in 96-well plate. Cell proliferation was determined by (H) anchorage-independent cell growth assay. The expression level of GAPDH was used as the control for Western blot. Cells incubated with secondary antibody were served as an isotype control for flow cytometry assay. Data is presented mean ± SEM values of three separate experiments in triplicate. \**P*<0.05, \*\**P*<0.01 and \*\*\**P*<0.001 indicate significance differences compared with untreated (control) group. The panels were magnified ×10 (Bar = 100µm).

**Figure 2.**
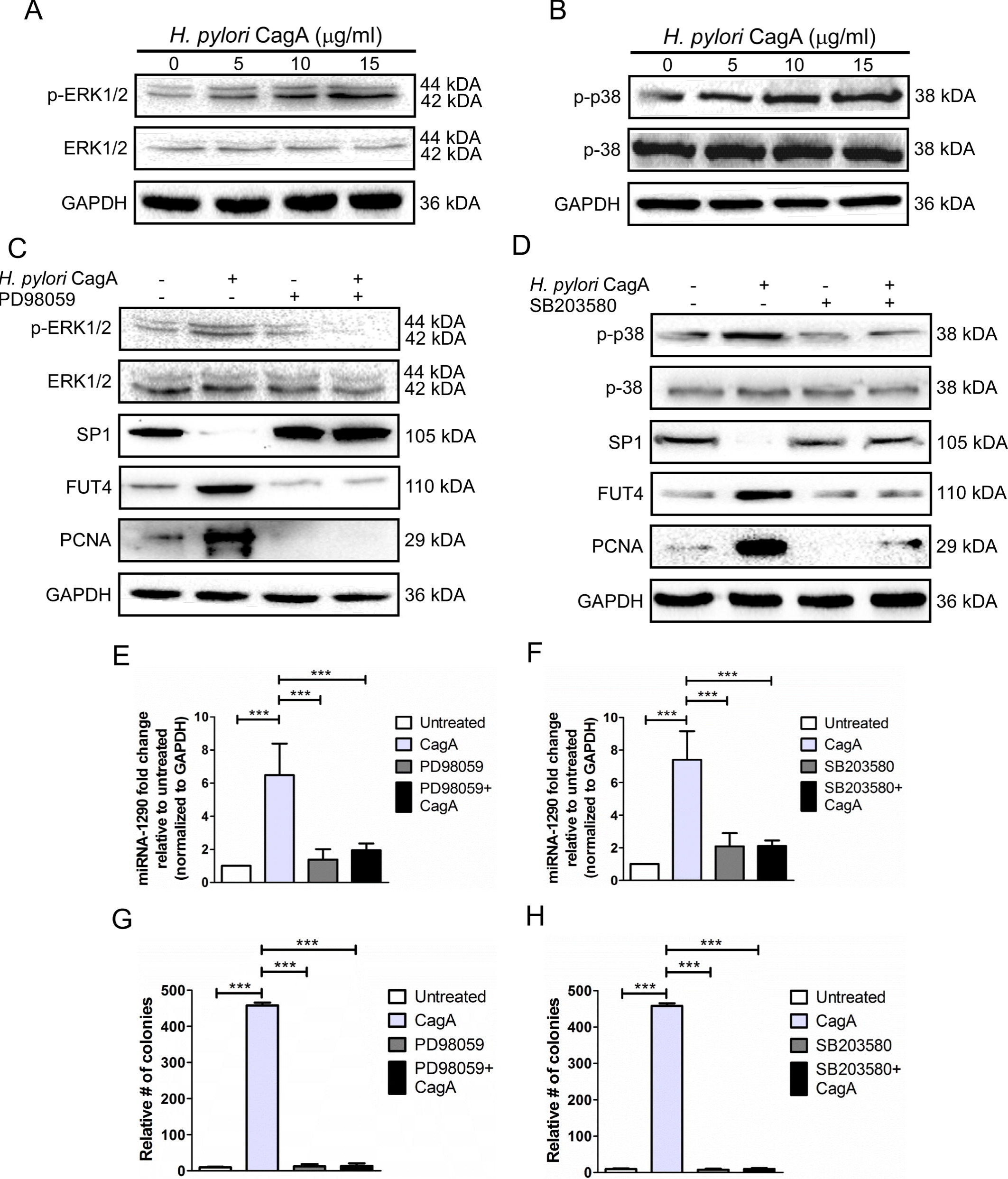
*H. pylori* CagA increases ERK1/2 and p38 MAP kinase activation. (A & B) p-ERK1/2 and p-P38 was detected by Western blot after treatment with CagA. (C & D) The human gastric cancer cells (SGC-7901) were treated with ERK1/2 (PD98059; 10^−5^ M) and p-38 (SB203580; 10^−6^ M) inhibitors for 24 h in serum free medium followed by CagA treatment. Total protein was isolated from the whole cell lysate and p-ERK1/2, p-P38 SP1, FUT4 and PCNA expression was detected by Western blot. (E & F) Relative miRNA-129 expression was determined by real time PCR after treatment of *H. pylori* CagA or ERK1/2 and p-38 inhibitors. The relative expression level of miRNA-1290 was determined by real time PCR after ERK1/2 and p38 inhibitors followed by CagA treatment. (G & H) Cell proliferation was determined by anchorage-independent cell growth assay. The expression level of GAPDH was used as the control for Western blot. Data is presented mean ± SEM values of three separate experiments in triplicate. \**P*<0.05, \*\**P*<0.01 and \*\*\**P*<0.001 indicate significance differences compared with untreated (control) group.

### H. pylori CagA activates miRNA-1290 and SP1 downregulation through MAPKs signaling pathway

The phosphorylation of ERK1/2 and P38 were examined to determine the role of EGFR/MAPKs signaling through miRNA-1290 activation in *H. pylori* CagA-induced cell proliferation. As shown in Figures 2A and 2B, *H. pylori* CagA treatment phosphorylated the ERK1/2 and P38 as compared to untreated cells. Furthermore, to identify the MAPKs signal pathway mechanism involved in *H. pylori* CagA associated up-regulation of miRNA-1290, cells were pretreated with ERK (10^−5^ M PD98059) and p38 inhibitors (10^−6^ M SB203580), followed by *H. pylori* CagA for 48 h. We found significant inhibition of *H. pylori* CagA-activated ERK1/2 and p38 phosphorylation, miRNA-1290, FUT4, as well as gastric cancer cell proliferation, while showed upregulation of SP1 (*P*<0.001) (Figures 2C, 2D, 2E. 2F). Moreover, We found *H. pylori* CagA exhibited a significantly stimulatory effect on human gastric cancer cell into colony transformation in a dose-dependent manner, while inhibited in ERK1/2 and p38 inhibitor treated cells (*P*<0.001) (Figures. 2G, 2H & Supplementary Figs. S2). These results suggest that ERK and p38 inhibitors blocked the stimulatory effect of *H. pylori* CagA on ERK1/2 and P38 activation respectively, leads to inhibition of miRNA-1290, FUT4 and CagA-induced cell proliferation with the stimulation of SP1 expression.

### Effect of H. pylori CagA on FUT4/LeY fucosylation through miRNA-1290 activation

Gastric cancer cells were treated with *H. pylori* CagA to analyse the effect of CagA on FUT4 fucosylation through miRNA-1290 activation. By real-time PCR (Figure 3A), flowcytometry assay, Western blot (Figure 3B, 3J), and ELISA (Figure 3C, 3D), we found significantly high expression of FUT4 and LeY level (*P*<0.001) in a dose-dependent manner in *H. pylori* CagA treated cells (Figure 3 & Supplementary Fig. S3). To understand the FUT4 fucosylation mechanism involved in *H*.*pylori* CagA-induced gastric cancer cell proliferation via miRNA-1290 activation, we analysed the inhibition of FUT4 by SiRNA knockdown assay and found a significant decreased expression of FUT4 (Figure 3E, 3F) and LeY level (Figure 3G, 3K) which leads to decreased gastric cancer cell proliferation in CagA treated cells, while SP1 showed significantly high expression as compared to CagA treated cells (*P*<0.001) (Figure 3H & Supplementary Fig. S3). However, the effect of *H. pylori* CagA on expression level of type I and type II Lewis antigens was analysed by Western blot and found significant effect of *H. pylori* CagA on LeB, LeX and sLeX as compared to LeA expression level (*P*<0.01) (supplementary Fig. S4). Furthermore, FUT4 SiRNA transfection was showed negligible difference of miRNA-1290 level in CagA treated cells (*P*>0.001) (Figure 4I). These results are probable indications that *H. pylori* CagA possibly stimulates the cell proliferation with up and down-regulation FUT4 fucosylation and SP1, respectively. However, FUT4 have no upstream role in the activation of miRNA-1290.

**Figure 3.**
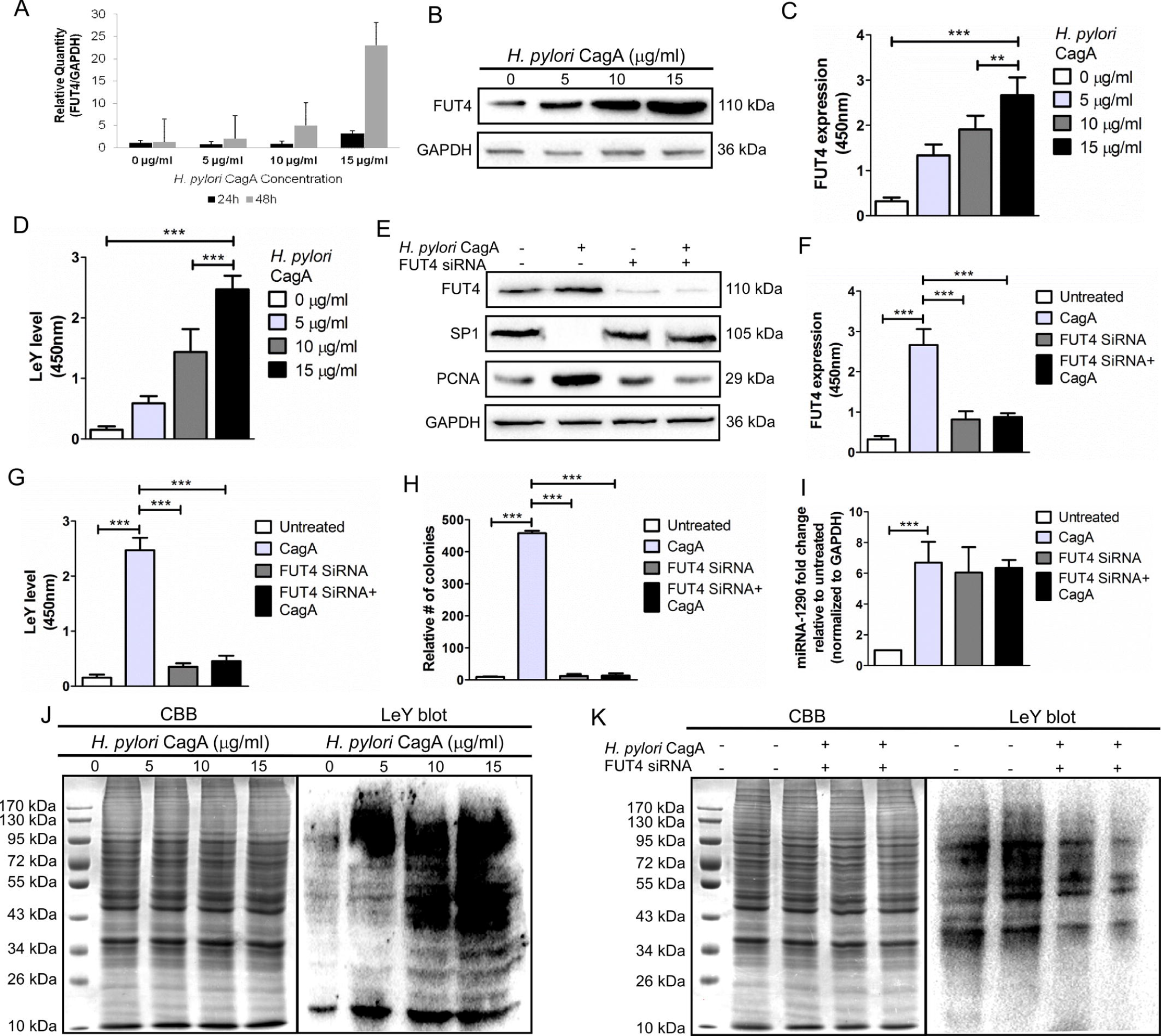
*H. pylori* CagA stimulates the FUT4-fucosylation and Lewis type I antigen (LeY) antigen in gastric cancer cells (SGC-7901). FUT4 and LeY expression level was detected in CagA treated cells by (A) real-time PCR, (B and J) Western blot, (C and D) ELISA. (E, K) Knocking down the expression of FUT4 decreased FUT4 and LeY expression and gastric cancer cell proliferation in the CagA overexpressed cells while SP1 expression was significantly increased. FUT4 and LeY expression was examined by (F and G) ELISA after FUT4 siRNA transfection followed by CagA treatment. (H) Human gastric cancer cells were treated with FUT4 siRNA (400 ng of plasmid). Cells (1 × 10^3^ cells/well) were seeded in 96-well plate and cell proliferation was determined by anchorage-independent cell growth assay. (I) miRNA-1290 level was determined by real time PCR after FUT4 SiRNA transfection. The expression level of GAPDH was used as the control for Western blot. Cells incubated with secondary antibody only served as an isotype control for flow cytometry assay. Data is presented mean ± SEM values of three separate experiments in triplicate. \**P*<0.05, \*\**P*<0.01 and \*\*\**P*<0.001 indicate significance differences compared with untreated (control) group.

**Figure 4:**
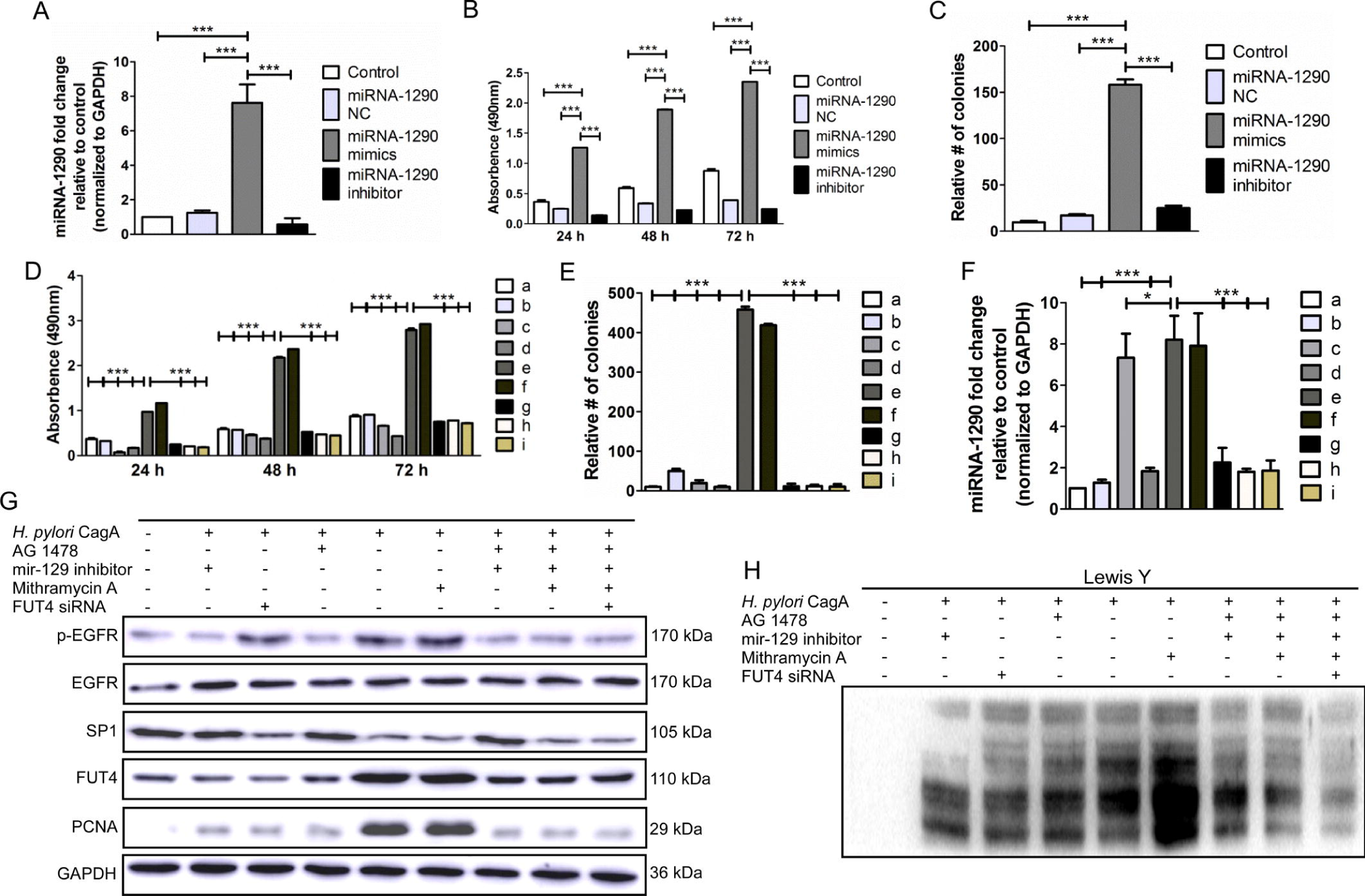
*H. pylori* CagA induced gastric cancer cell proliferation by regulating miRNA-1290. (A) The relative expression level of miRNA-1290 was measured in SG7901 cells transfected with miRNA-1290 mimics and inhibitor by real time RCR, (B and C) as well cell proliferation by MTT and anchorage-independent cell growth assay. (D and E) Human gastric cancer cells were treated with inhibitor of EGFR inhibitor Tyrphostin AG 1478 (10^−4^ M), SP1 (10^−10^ M Mithramycin A), miRNA-1290 (10^−6^ M KRIBB11) and FUT4 siRNA (400 ng of plasmid), followed by CagA (15 µg/ml) for 48h. Cells (1 × 10^3^ cells/well) were seeded in 96-well plate. Cell proliferation was determined by MTT proliferation and anchorage-independent cell growth assay. (F) miRNA-1290 level was determined by real time PCR. (G and H) The expressions of pEGFR, SP1, FUT4, cell proliferation (PCNA) and LeY level were detected by Western blotting. (a) untreated, (b) miRNA-1290, (c) FUT4 siRNA EGFR, (f) SP1, (g) EGFR + miRNA-1290, (h) EGFR+ miRNA-1290 + SP1, (i) EGFR + miRNA-1290, (j) EGFR + miRNA-1290 + SP1 + FUT4 siRNA, followed by CagA (15 µg/ml) for 48h. The expression level of GAPDH was used as the control for Western blot. Data is presented mean ± SEM values of three separate experiments in triplicate. \**P*<0.05, \*\**P*<0.01 and \*\*\**P*<0.001 indicate significance differences compared with (a) untreated (control) group.

### miRNA-1290 induces H. pylori treated gastric cancer cell proliferation through EGFR/SP1/FUT4/LeY pathway

In order to understand the mechanisms resulting in the promotion of gastric cancer cell proliferation through miRNA-1290 activation, SGC7901 cells were transfected with miRNA-1290 mimics; showed significant high cell proliferation and miRNA-1290 expression as compared to inhibitor and negative control (Figure 4A, 4B, 4C & Supplementary Fig. S5) (*P*<0.001). The miRNA-1290 inhibition experiments in gastric cancer cell confirmed our hypothesis that miRNA-1290 targeted the SP1 and FUT4 genes which lead to proliferate gastric cancer (Figure: 4). Real-time PCR, Western blot, and cell proliferation (CCK8) and anchorage-independent cell transformation assay results (4F and Supplementary Fig. S5) indicated that miRNA-1290 deactivation with EGFR inhibition significantly decreased the FUT4 (Figure 4G), LeY (Figures 4H) and PCNA expression (*P*<0.001), whereas SP1 expression increased to prevent the development of cell proliferation (*P*<0.001). In addition, inhibition of SP1 (10^−10^ M Mithramycin A) and FUT4 (FUT4 siRNA) lead to significantly up- and down-regulation of gastric cancer cell proliferation, respectively (Figures 4D, 4E & Supplementary Fig. S5) (*P*<0.001). However, we were not found upstream effect of SP1 and FUT4 inhibition on miRNA-1290 (*P*<0.001) (Figures 4F). These data correlate well with our finding that overexpression of miRNA-1290 by *H. pylori* CagA caused increased in gastric cancer cell proliferation through EGFR/MAPKs/SP1/FUT4/LeY signaling pathway (Figure. 6).

**Figure 5:**
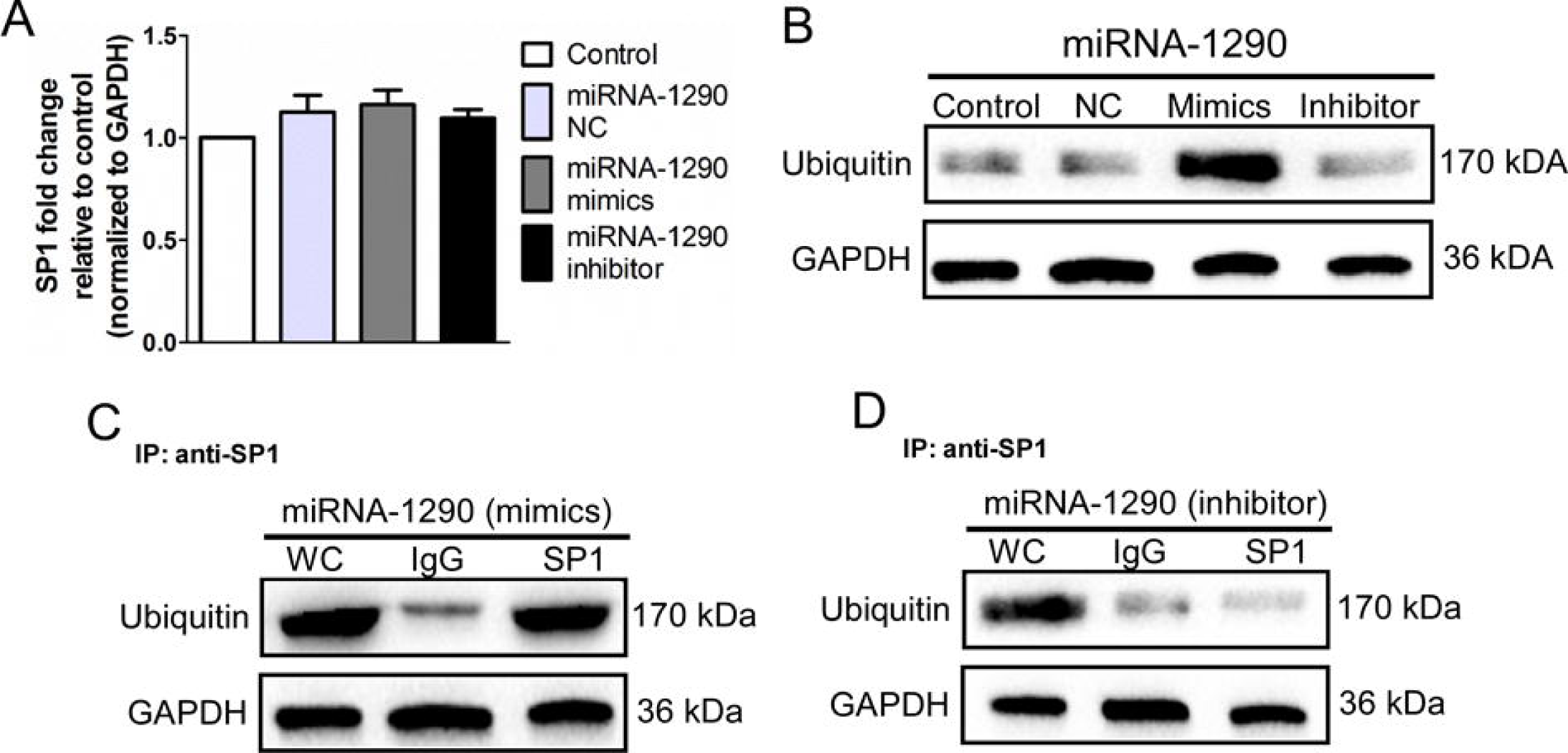
miRNA-1290 regulated SP1 expression through ubiquitination proteasomal interaction. *H. pylori* CagA induced gastric cancer cell proliferation by regulating miRNA-1290. (A) The mRNA level of SP1 was determined in SG7901 cells transfected with miRNA-1290 inhibitor by real time RCR. (B) The expression level of ubiquitin was measured in SG7901 cells transfected with miRNA-1290 mimics and inhibitor by western blot. (C and D) Cell lysates extracted were utilized for immunoprecipitation using an anti-SP1 antibody with mimics and inhibitor of miRNA-1290; the immune complexes were analyzed by Western blotting with ubiquitin antibodies. The expression level of GAPDH was used as the control for Western blot. Data is presented mean ± SEM values of three separate experiments in triplicate. \**P*<0.05, \*\**P*<0.01 and \*\*\**P*<0.001 indicate significance differences compared with (a) untreated (control) group.

**Figure 6:**
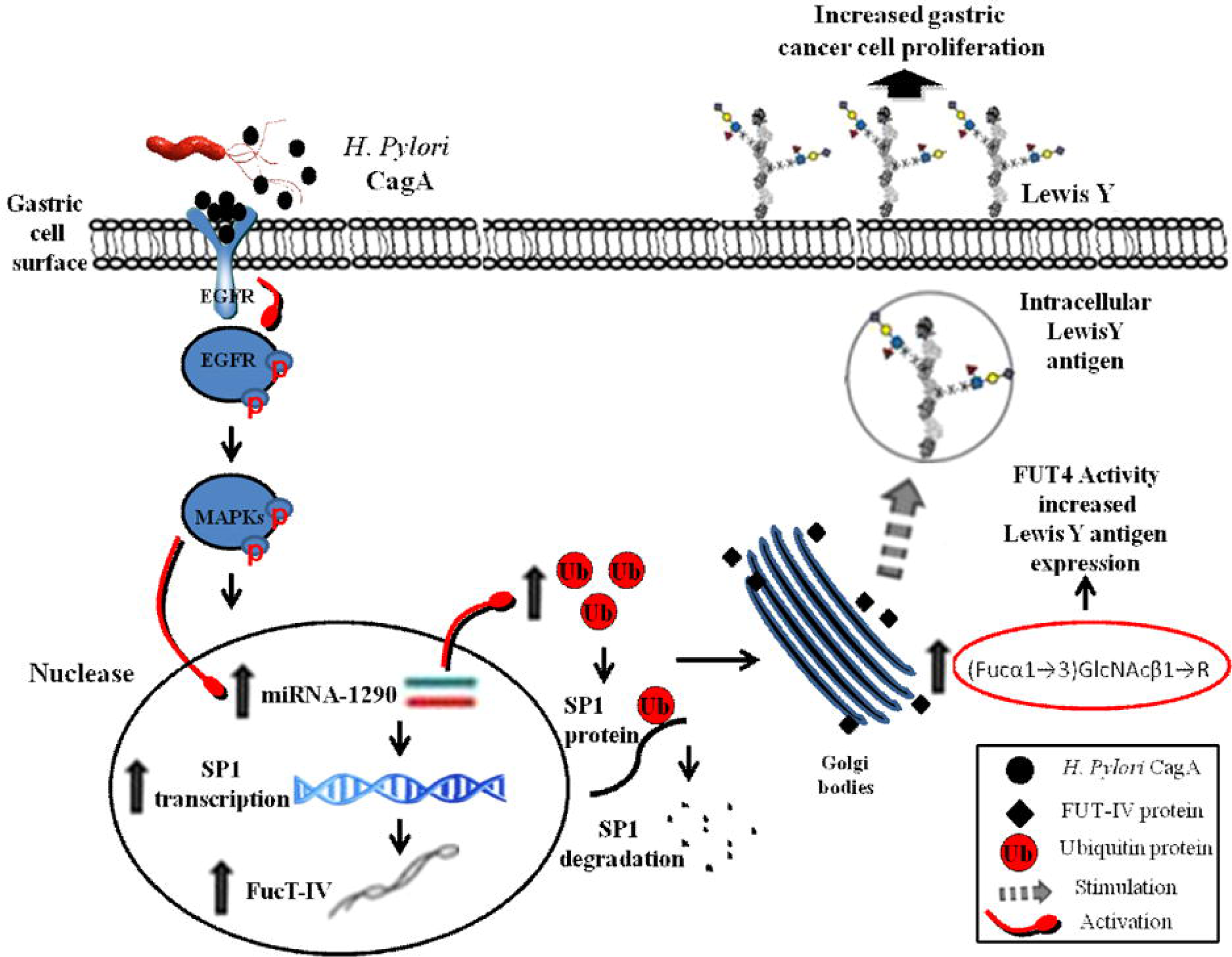
Schematic representation of *H. pylori* CagA associated gastric cancer cell proliferation mechanism through miRNA-1290 regulation with overexpression of FUT4/LeY. *H. pylori* regulated the miRNA-1290 through EGFR/MAPKs phosphorylation, which overexpressed the FUT4/LeY level with high gastric cancer cell proliferation. However, degradation of SP1 transcriptional activity by ubiquitin proteasomal system increases the biosynthesis of FUT4 and stimulates the gastric cancer cell proliferation.

### miRNA-1290 inhibits SP1 expression through ubiquitination proteasomal pathway

In order to analyze the mechanism responsible for the degradation of SP1 protein, we examined the expression level of SP1 in miRNA-1290 treated cells, and we found negligible differences of SP1 mRNA level as compared to control (Figure. 5A); which proposed that SP1 is degraded in protein phase rather at mRNA level. We examined the potential role of ubiquitin-proteasomal pathway and found significantly high expression of ubiquitin protein level in response to miRNA-1290 treatment (*P*<0.001) (Figure 5B). Later, the direct interaction between sp1 and ubiquitin was determined by immunoprecipitation. As shown in Figure 5, the role of ubiquitin activation showed that SP1 directly interact with ubiquitin-proteasomal system as we found significantly high expression of ubiquitin in miRNA mimics treated cells, as compared to miRNA-1290 inhibitor treated cells (Figure. 5C, 5D). These results predict the likely involvement of ubiquitin protein in the proteasomal degradation of SP1 protein with the activation of miRNA-1290 in *H. pylori* CagA treated gastric cancer cells.

## Discussion

In recent years, the role of miRNAs in pathogenesis of gastric cancers has been extensively studied. However, little is known regarding on the association between miRNAs and cancer glycosylation responses in *H*.*pylori*-associated gastric cancer. Recently, our lab was found a critical role of mir200b in the inhibition of breast cancer proliferation and metastasis by targeting fucosyltransferase IV and α1,3-fucosylated glycans [37]. Previously, we reported that Rg3 drug stimulated gastric cancer cell apoptosis through FUT4 fucosylation by regulating SP1 and HSF1 expressions, which suggested oncogenic role of CagA in development of *H. pylori* associated gastric cancer [34], [35]. Here, we investigated the detail mechanism of miRNA-1290 to promote gastric cancer cell proliferation by targeting SP1 through fucosyltransferase IV and α1,3-fucosylated glycans. *H. pylori* CagA directly or indirectly induces various molecular changes in gastric epithelial cells, which facilitate the cell proliferation [38]. Previously, we reported that inhibition of EGFR phosphorylation downregulated gastric cancer cell proliferation through MAPKs pathways [39]. Hence, we evaluated the role of CagA on downstream molecules of EGFR and miR-1290 through MAPKs pathway. Our results showed that phosphorylation of EGFR were associated with up and down-regulation of miRNA-1290 and SP1 through ERK1/2 and p38 activation. According to Keates et al., *H. pylori* CagA positive strain was more potent to induce stronger MAPKs phosphorylation of ERK and p38 in AGS cell line [40]. These results proposed that miRNA-1290 is activated through EGFR/MAPKs phosphorylation by *H. pylori* CagA. Hence, *H. pylori* CagA is a microbial factor associated with developmental risk of gastric cancer and its expression may have potential application as a marker for the evaluation of this risk.

*H. pylori* infection leads to the abnormal glycosylation and induced expression of the specific FUTs in gastric mucosa [41], [42], [43]. *H. pylori* can recognize a series of host glycosylated molecules, including LeY antigen which is predominately expressed by the mucous, chief and parietal cells of gastric glands [11]. Faisal et al., reported high expression of FUT4 and LeY in gastric cancer (97 & 92%) as compared to gastritis (82% & 85%) and gastric ulcer (82% & 85%) [35]. In this study, we found that *H. pylori* CagA is associated with high expression of FUT4 and LeY in gastric cancer cells with dose dependent manner. Furthermore, FUT4 inhibition proved that FUT4 fucosylation and α1,3-fucosylated glycans (LeY) are responsible for gastric cancer cell proliferation (Figure 4). However, miRNA-1290 expression was not inhibited with the downregulation of FUT4, which suggests that miRNA-1290 is not be affected by FUT4 inhibition and placed at upstream of FUT4 and LeY (Figures 6). FUT4 overexpression promotes cell proliferation through elevating LeY synthesis in human epidermoid carcinoma [15]. Faisal et al., investigated the therapeutic role of anti-LeY antibody to enhance the efficacy of celecoxib against gastric cancer by downregulating MAPKs/COX-2 signaling pathway [3]. Hence, this results suggests that miRNA-1290 is not directly associated with FUT4 fucosylation and α1,3-fucosylated glycans (LeY). However, some other proteins might play critical role through miR-1290 to promote fucosylation in gastric cancer.

*H. pylori* can recognize a series of host glycosylated molecules for adhesion and infection by their adhesins. In human stomach, the LeY blood antigen is predominately expressed by the mucous, chief and parietal cells of gastric glands [44]. Hence, we also investigated the role of CagA on Type I [Gal-β (1,3)-GlcNAc] and II [Gal-β (1,3)-GlcNAc] human Lewis sugar antigens; our results indicated that *H. pylori* CagA can modify the intensity of the gastric Lewis antigen level in gastric cancer; while Lewis Y was found significantly upregulated as compared to LeA(1−4 location), LeB (1−2 and 1−4), LeX (1−3 position), sLeX (on both 2−2 and 2−4 location). *H. pylori* expresses BabA and SabA adhesins to facilitate its colonization with the gastric epithelium surface by the host LeB, LeX and sLeX antigens [11], [45]. According to Wirth et al., high expression level of host Lewis antigen might be particularly adaptive in CagA strain of *H*.*pylori* [46]. Hence, there is a need to further investigate and analysed the role of *H. pylori* CagA on host LeA, LeB, LeX and sLeX antigen in correlation with their respected FUTs enzyme.

The detail mechanism of miRNA-1290 in the promotion of gastric cancer through FUT4/LeY fucosylation was further confirmed by using inhibition assay experiment. We hypothesized that *H. pylori* CagA owned an ability to up-regulate miRNA-1290, due to phosphorylation of EGFR/MAPKs and leads to inhibit SP1 protein with the regulation of ubiquitin proteasomal system. In detail, miRNA-1290 inhibition through EGFR inhibitor (AG-1478) leads to suppress gastric cancer cell proliferation. In contrast, miRNA-1290 activation through SP1 knockdown proposed that miRNA-1290 has significant association with SP1 protein degradation. Furthermore, loss of SP1 expression in *H. pylori*-associated gastric cancer was found due to the activation of ubiquitin proteasomal pathway and which promotes gastric fucosylation and gastric cancer cell proliferation. SP1 protein degradation by ubiquitin proteasomal system was investigated in an effort to understand the SP1 damage at protein level induced by the miRNA-1290. The ubiquitin-proteasome system is an important cellular protein degradation pathway. E2, E3, deubiquitination enzymes (DUBs) and proteasomes play an important role in the tumorinegenesis and cancer development [47]. Our result proposed that activation of ubiquitin proteasome system leads to stimulate FUT4/LeY fucosylation and promotes gastric cancer cell proliferation by degrading SP1 protein. Further investigation carried out by imunoprecipitation results, suggested that ubiquitin protein is directly associated with the SP1 inhibition with the regulation of miR-1290. Hence, tumor suppressor transcription factor activity of SP1 is thought to negatively regulate the miRNA-1290 and followed to inhibit gastric cancer cell proliferation.

## Conclusion

Taken together, it is tempting to reveal that *H. pylori* CagA stimulates miRNA-1290 expression which target the SP1 protein expression in association with ubiquitin proteasomal system and facilitates the overexpression of FUT4 fucosylation and α1,3-fucosylated glycans in the gastric epithelium (Figure 6). This study provide a new interconnection between miRNA-1290 and gastric cancer fucosylation. Hence, miRNA-1290 can serve as promising target for interventional strategies towards gastric cancer treatment.

## Abbreviations

CagA: Cytotoxin associated antigen A
EGFR: epidermal growth factor receptor
ERK1/2: extracellular signal-regulated kinases
FACS: fluorescence activated cell sorting
FUTs: fucosyltransferases
ICC: immunocytochemistry
IFC: immunofluorescence
LeA: Lewis A
LeB: Lewis B
LeX: Lewis X
LeY: Lewis Y
MAPKs: mitogen-activated protein kinases
miRNA: microRNA
PCNA: proliferation cell nuclear antigen
p-EGFR: phosphorylated epidermal growth factor receptor
sLeX: sialyl-Lewis X
SP1: specificity protein 1

## Acknowledgments

The project was supported by the China 973 grant no. 2012CB822100, National Natural Science Foundation of China Research grants no. 30672753 and 31270866.

## Figure legends

**Supplementary Figure S1:** SG7901 cancer cells were treated with different concentration of *H. pylori* CagA (15 µg/ml) and assayed for their ability to proliferate through EGFR activation. The expression of (A) PCNA and (B) EGFR was detected of post CagA treatment by (A) immune-cytometry and (B) flow cytometry assay. (C and D) MTT and (E) soft agar assays were performed of SG7901 cancer cells and were treated with Tyrphostin AG 1478 (10^−4^ M) for 24 h in serum free medium followed by *H. pylori* CagA (15 µg/ml) and assayed for their ability to proliferate. Cells incubated with secondary antibody were served as an isotype control for flow cytometry assay. Data is presented mean ± SEM values of three separate experiments in triplicate. \**P*<0.05, \*\**P*<0.01 and \*\*\**P*<0.001 indicate significance differences compared with untreated (control) group. The panels were magnified ×10 (Bar = 100µm).

**Supplementary Figure S2:** *H. pylori* CagA stimulates anchorage-independent soft agar growth of gastric cancer cell through activation of ERK1/2 and P38. SG7901 cancer cells were treated with ERK1/2 (PD98059; 10^−5^ M) and P38 (SB203580; 10^−6^ M inhibitors for 24 h in serum free medium followed by *H. pylori* CagA (15 µg/ml) and assayed for their ability to proliferate by MTT and soft agar. Multicellular colony formation was photographed at 10× magnification. Data is represented as mean ± SEM (n = 3). \**P*<0.05, \*\**P*<0.01 and \*\*\**P*<0.001 vs untreated group).

**Supplementary Figure S3:** The expression of (A) FUT4 and (B) LeY was detected of post CagA treatment by (A and B) flow cytometry assay. SG7901 cancer cells were treated with FUT4 siRNA (400 ng of plasmid) for 24 h in serum free medium followed by *H. pylori* CagA (15 µg/ml) and assayed for their ability to proliferate by MTT and soft agar. Cells incubated with secondary antibody were served as an isotype control for flow cytometry assay. Data is presented mean ± SEM values of three separate experiments in triplicate. \**P*<0.05, \*\**P*<0.01 and \*\*\**P*<0.001 indicate significance differences compared with untreated (control) group. The panels were magnified ×10 (Bar = 100µm).

**Supplementary Figure S4:**

*H. pylori* CagA treatment up regulates the Lewis type I and type II antigen level in human gastric cancer cells (SGC-7901). Gastric cancer cells were treated with different concentrations of CagA (0–15 µg/ml) for 48 h. Total protein from the whole cell lysate was subjected to SDS–PAGE and blotted with respected antibodies. Effect of CagA on type I and type II sugar antigen was detected by western blot as (A) LeA, (B) LeB, (C) LeX, (D) sLeX. Coomassie Brilliant Blue (CBB) staining of gels showed comparable amounts of proteins in each lane (left panel). Data presented here were representative of mean values from three separate experiments in triplicate. \**P* < 0.05 and \*\**P* < 0.01 indicated significance differences compared with untreated (0 µg/ml) control group.

**Supplementary Figure S5:** *H. pylori* CagA stimulates anchorage-independent soft agar growth of gastric cancer cell through miRNA-1290 activation. SG7901 cancer cells were transfected with miRNA-1290 mimics and inhibitor in serum free medium followed by *H. pylori* CagA (15 µg/ml) and assayed cell proliferation by soft agar. Moreover, SG7901 cancer cells were treated with the inhibitor of (a) untreated, (b) miRNA-1290, (c) FUT4 siRNA (d) EGFR, (f) SP1, (g) EGFR + miRNA-1290, (h) EGFR+ miRNA-1290 + SP1, (i) EGFR + miRNA-1290, (j) EGFR + miRNA-1290 + SP1 + FUT4 siRNA, followed by (e) CagA (15 µg/ml) for 48h and assayed for their ability to proliferate in soft agar. Multicellular colony formation was photographed at 10× magnification. Data is represented as mean ± SEM (n = 3). \**P*<0.05, \*\**P*<0.01 and \*\*\**P*<0.001 vs (A) untreated group).

**Supplemental Table1:**

Conditions and primers of routine RT-PCR and Real-time PCR (qPCR)

