## supplementary figures for "miR-1290 stimulates proliferation of gastric cancer by targeting SP1 with overexpression of fucosyltransferase IV and α1, 3-fucosylated glycans"

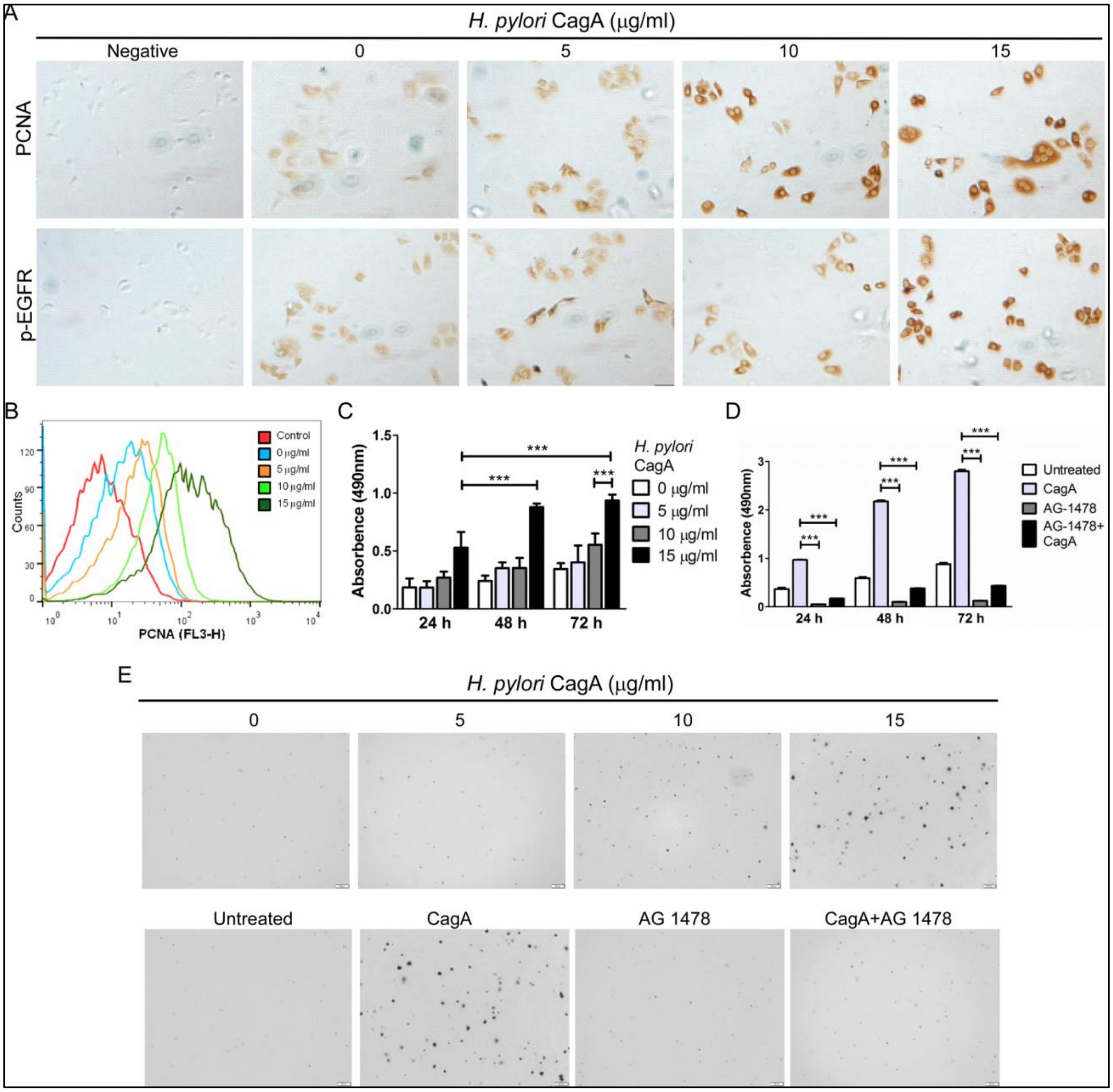


S1: Supplementary Figure 1

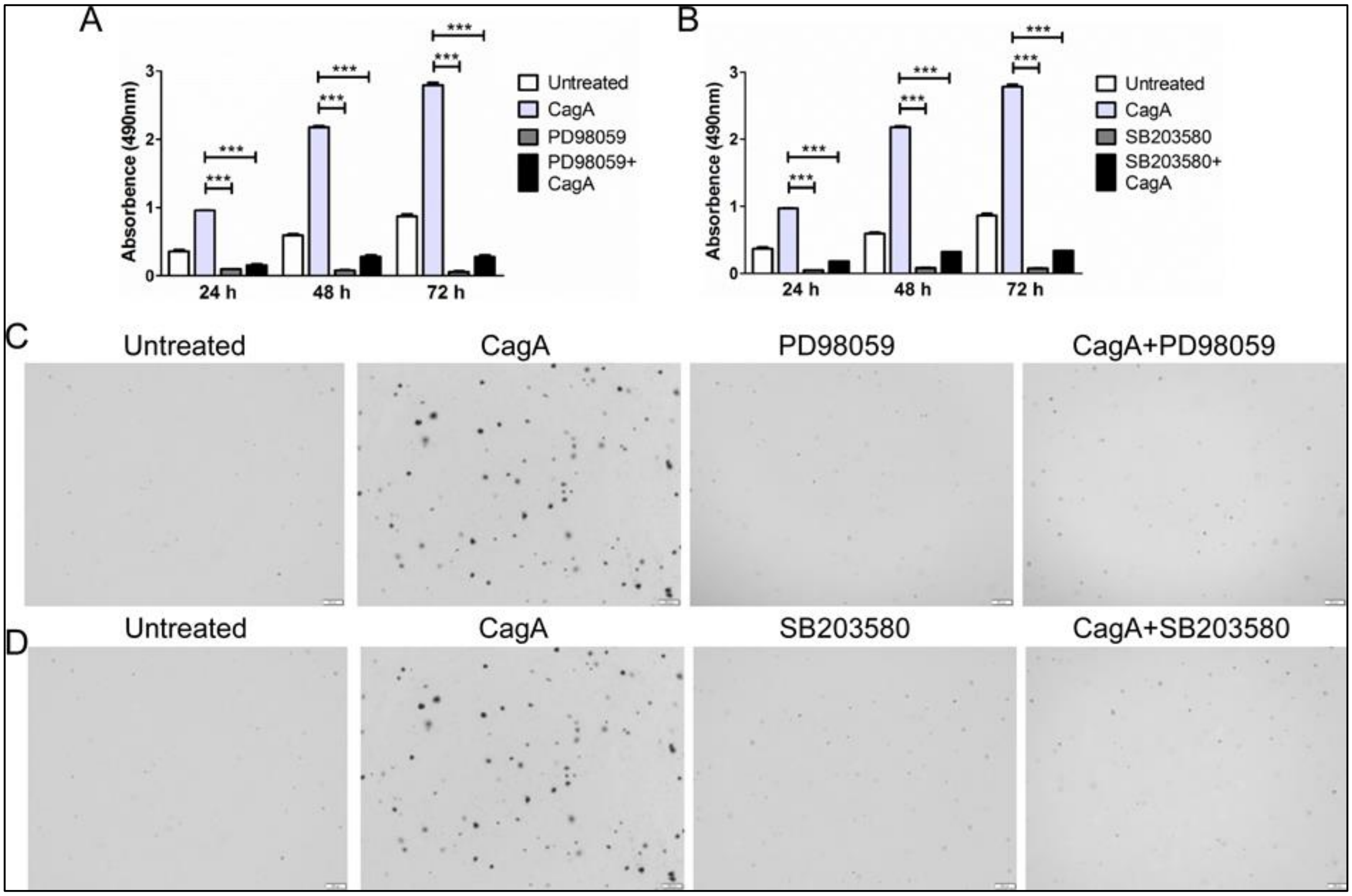


S2: Supplementary Figure 2

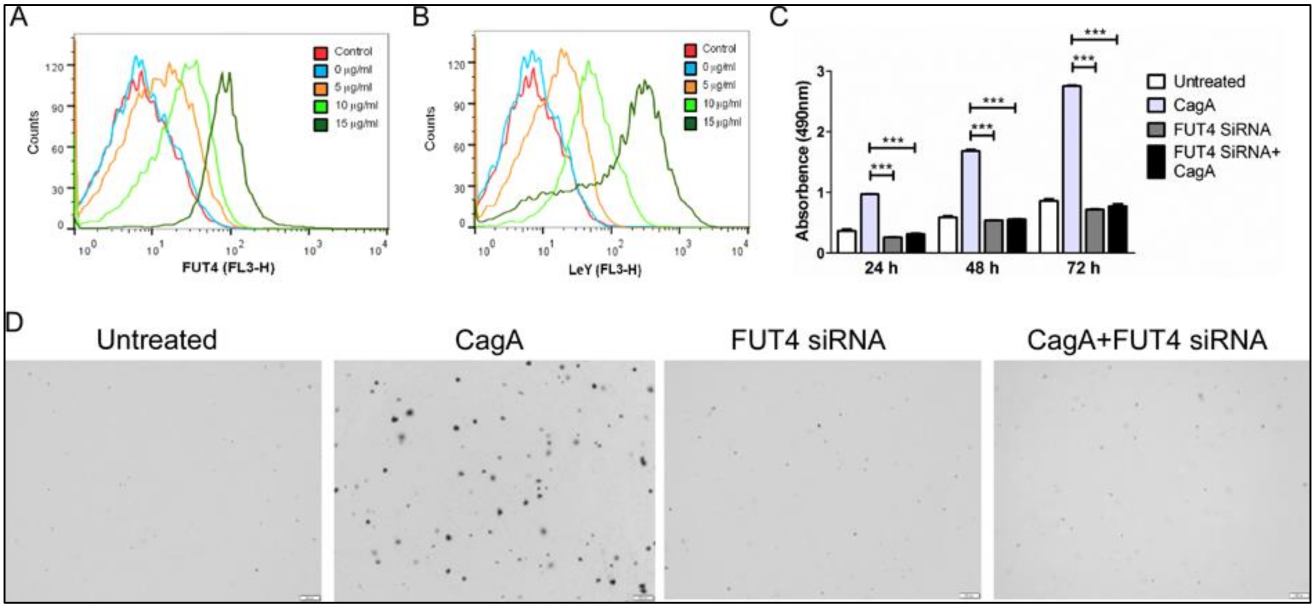


S3: Supplementary Figure 3

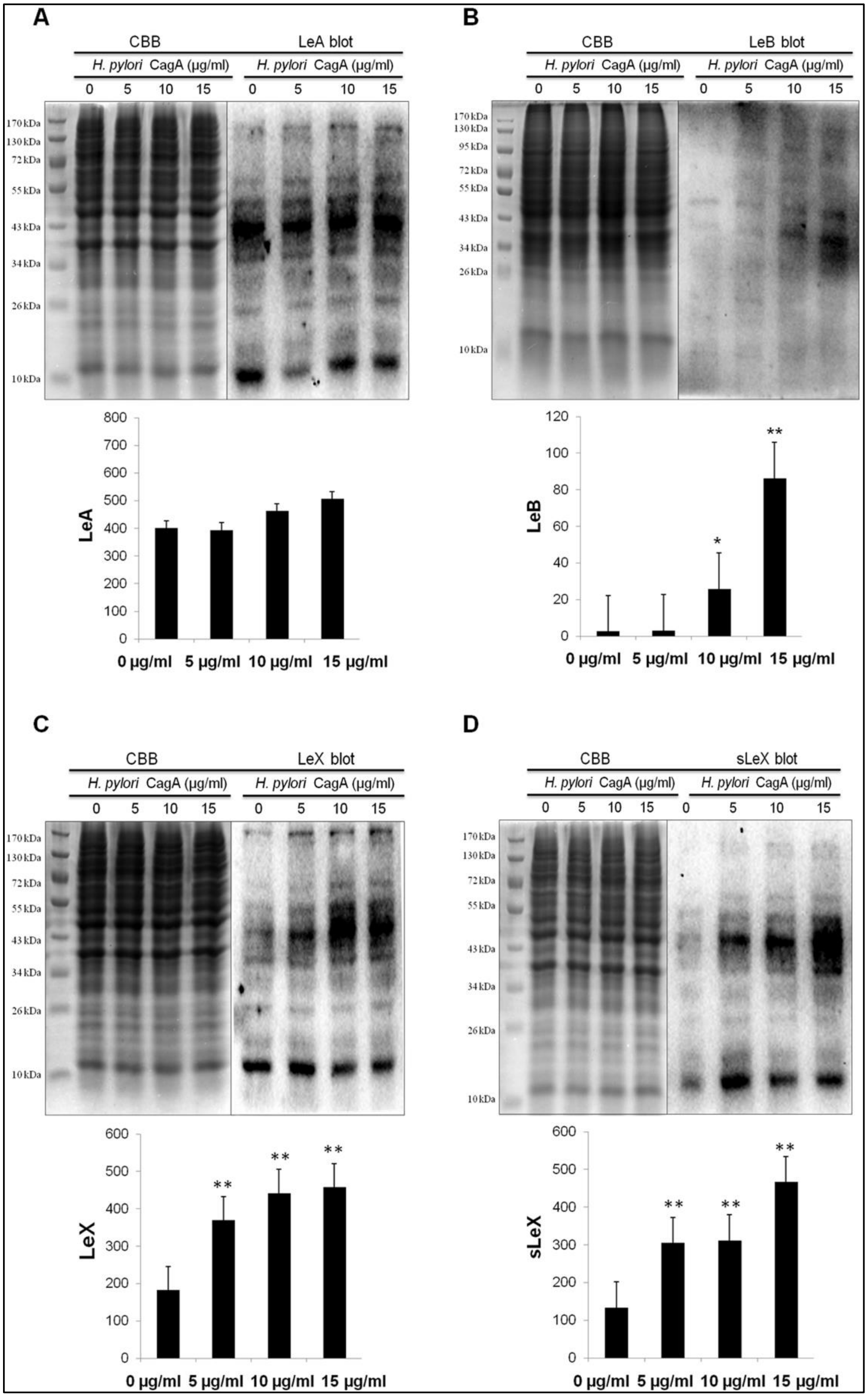


S4: Supplementary Figure 4

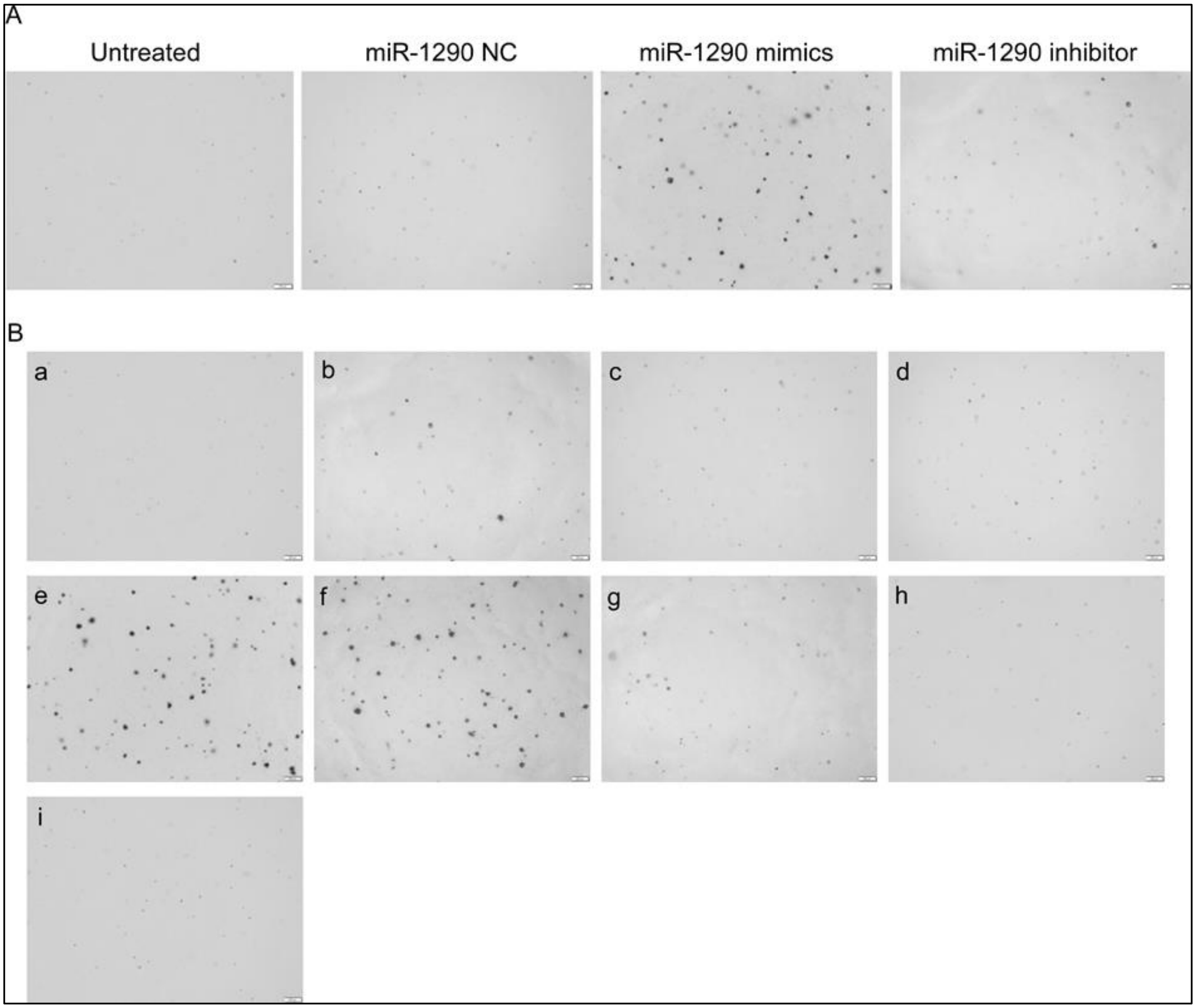


S5: Supplementary Figure 5
