## Supplementary table for "miR-1290 stimulates proliferation of gastric cancer by targeting SP1 with overexpression of fucosyltransferase IV and α1, 3-fucosylated glycans"

Supplemental Table1: Conditions and primers of routine RT-PCR and real-time PCR

| **Gene Name** | **Denaturation** | **Cycles** | **Extension** | **Primers** |
| --- | --- | --- | --- | --- |
| PCNA  (RT-PCR) | 94 ºC for 5 min | 28 cycles:  94 ºC for 30 s,  55 ºC for 30 s,  72 ºC for 30 s | 72 ºC for 10 min. | F:5’-TCAGCCTGTATCAAAGTCACC-3’ R:5’-ATCAGGAGCAGGAAAGCAAAG-3’ |
| GAPDH  (RT-PCR) | 94 ºC for 5 min | 28 cycles:  94 ºC for 30 s,  55 ºC for 30 s,  72 ºC for 30 s | 72 ºC for 10 min. | F:5’-ATGGGGAAGGTGAAGGTCG-3’ R: 5’-GGGGTCATTGATGGCAACAATA-3’ |
| FUT4  (qPCR) | 95 ºC for 30 s | 45 cycles:  95 ºC for 5 s,  60 ºC for 10 s, |  | F:5’-CTCAGGCCGTGCTTTTCCA-3’ R:5’-GTAGTCCAACACGCGCAGAT-3’ |
| miRNA-1290  (qPCR) | 95 ºC for 30 s | 45 cycles:  95 ºC for 5 s,  60 ºC for 10 s, |  | F:5’-GGAGTCCACTGGCGTCTTCAC-3’ R:5’-GAGGGGCCATCCACAGTCTTCT-3’ |
| SP1  (qPCR) | 95 ºC for 30 s | 45 cycles:  95 ºC for 5 s,  60 ºC for 10 s, |  | F:5’-GGAGTCCACTGGCGTCTTCAC-3’ R:5’-GAGGGGCCATCCACAGTCTTCT-3’ |
| GAPDH  (qPCR) | 95 ºC for 30 s | 45 cycles:  95 ºC for 5 s,  60 ºC for 10 s, |  | F:5’-GGAGTCCACTGGCGTCTTCAC-3’  R:5’-GAGGGGCCATCCACAGTCTTCT-3’ |
